## Supplementary Figures for "Functional evaluation of transposable elements as transcriptional enhancers in mouse embryonic and trophoblast stem cells"

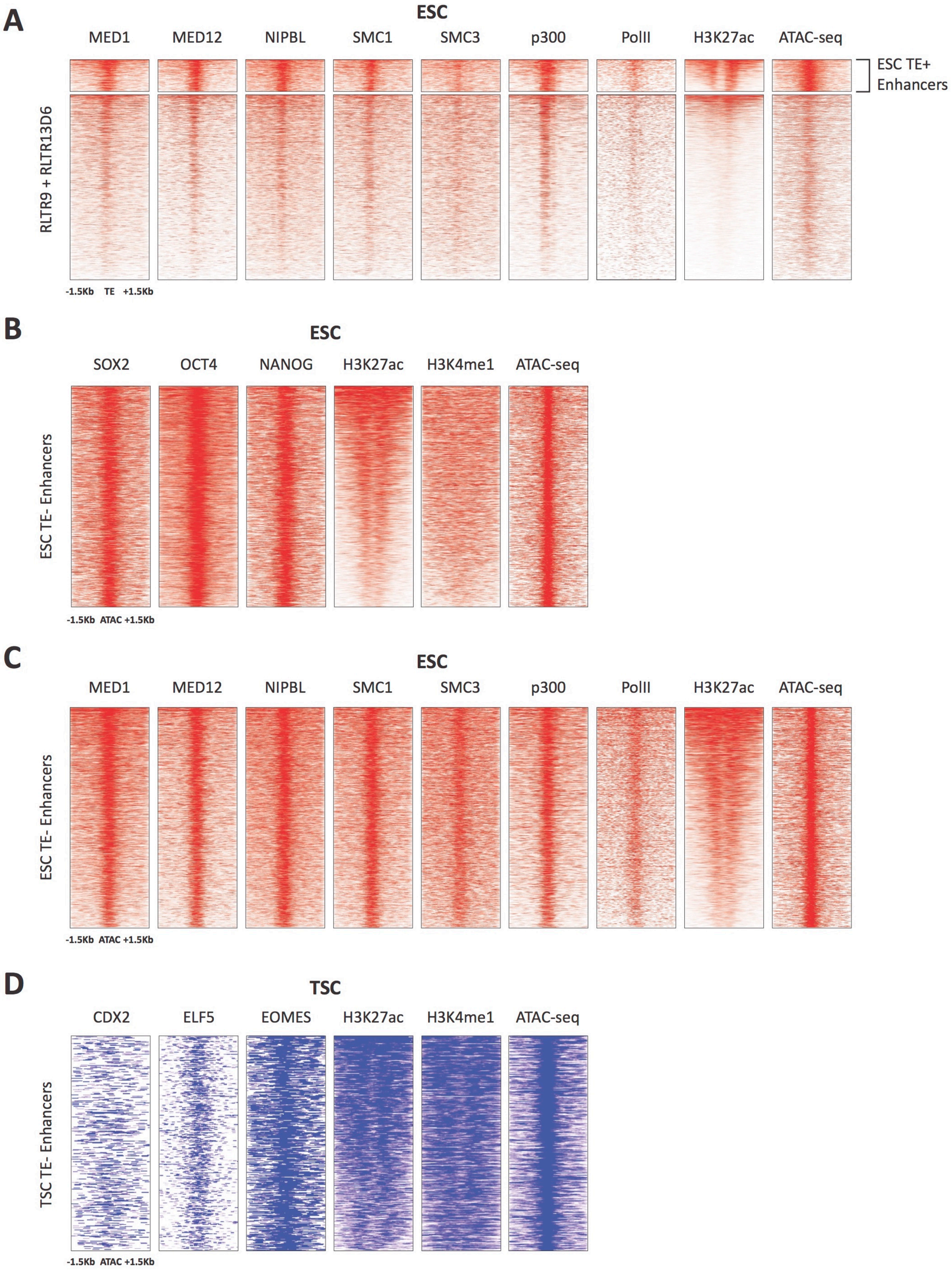


**Supplementary Figure 1 – Additional chromatin profiles of TEs and enhancers.** A) RLTR13D6 and RLTR9 ChIP-seq profiles for Mediator and cohesin complex components, p300 and RNA polymerase II, which are commonly associated with active enhancers. B) ChIP-seq and ATAC-seq profiles of ESC TE- enhancers, using the same data as in Figure 1B. C) Additional ChIP-seq profiles of TE-enhancers, using the same data as in panel A. D) ChIP-seq and ATAC-seq profiles of TSC TE-enhancers, using the same data as in Figure 1B.

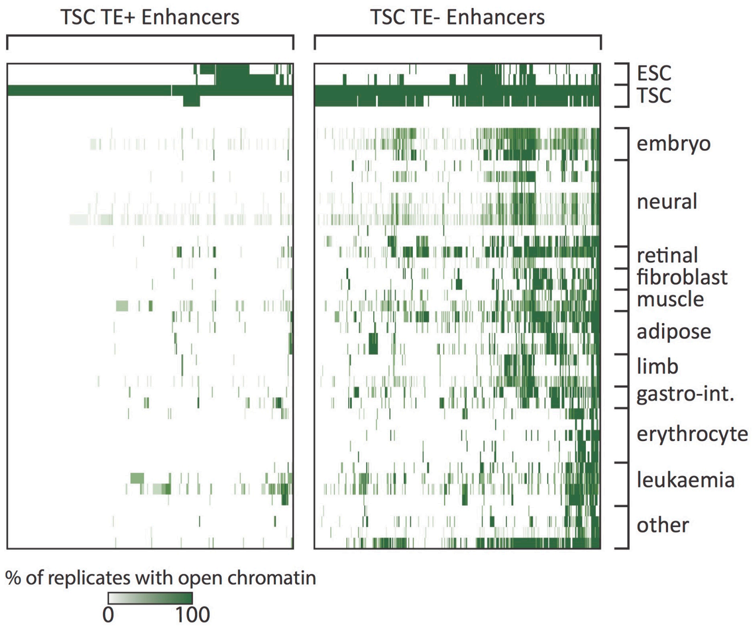


**Supplementary Figure 2 – Tissue specificity of TSC TE+ enhancers.** Analysis of ENCODE DNase-seq data from multiple tissues, displaying overlaps of open chromatin regions with TE+ and TE- enhancers in TSCs. For each TE, the colour intensity is proportional to the percentage of replicates of the same tissue that overlapped an open chromatin region.

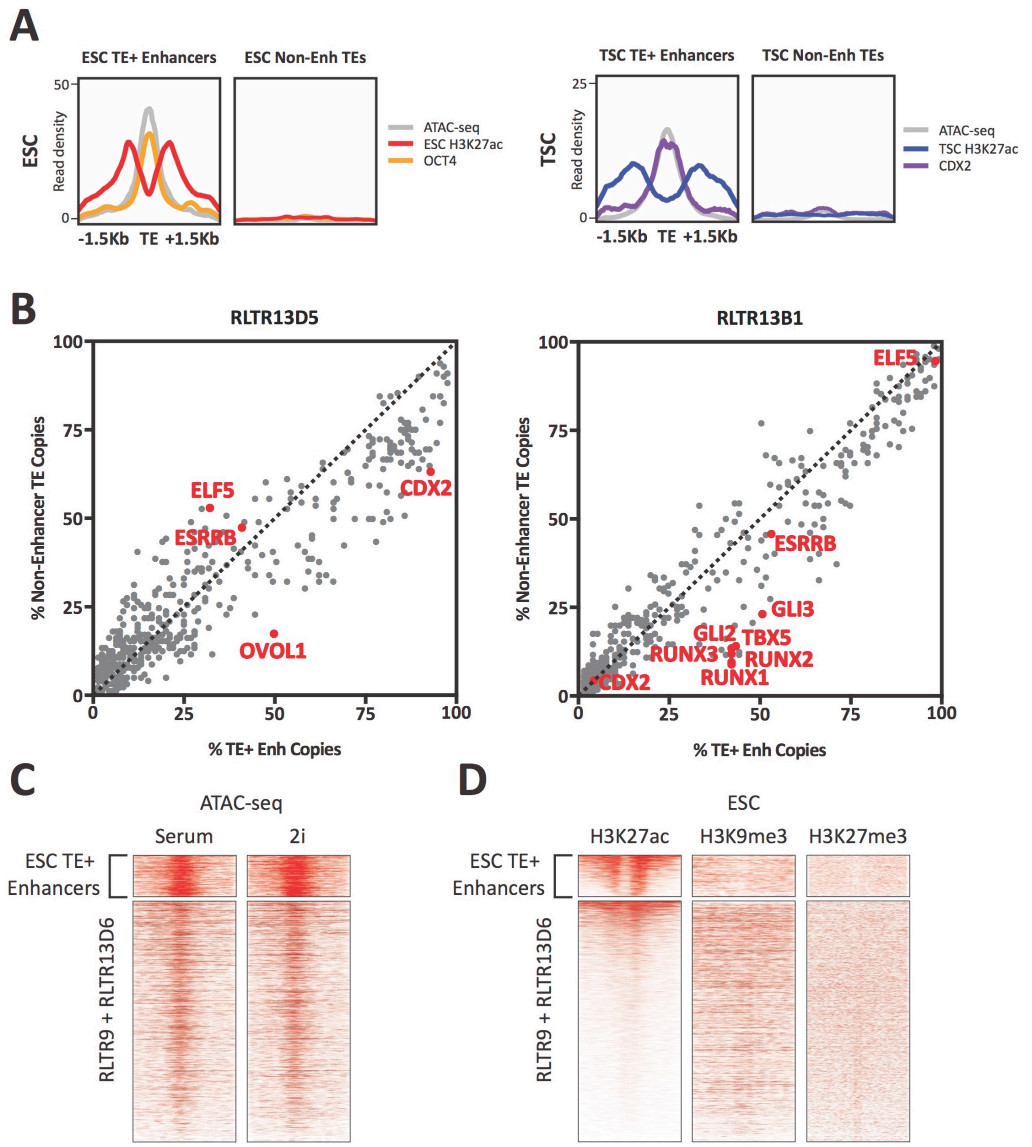


**Supplementary Figure 3 – Additional chromatin profiles of TEs and enhancers.** A) Average ChIP-seq and ATAC-seq profiles of selected non-enhancer TEs, compared with TE+ enhancers of the same subfamilies. B) Abundance of TF motifs at TSC TE+ enhancers and non-enhancer TEs from RLTR13D5 and RLTR13B1 subfamilies. C) ATAC-seq profiles of RLTR13D6 and RLTR9 elements in serum- or 2i-grown ESCs, showing similar profiles between both cell culture conditions. D) ChIP-seq profiles of RLTR13D6 and RLTR9 elements for H3K9me3 and H3K27me3, suggesting that non-enhancer TEs are not repressed by these marks.

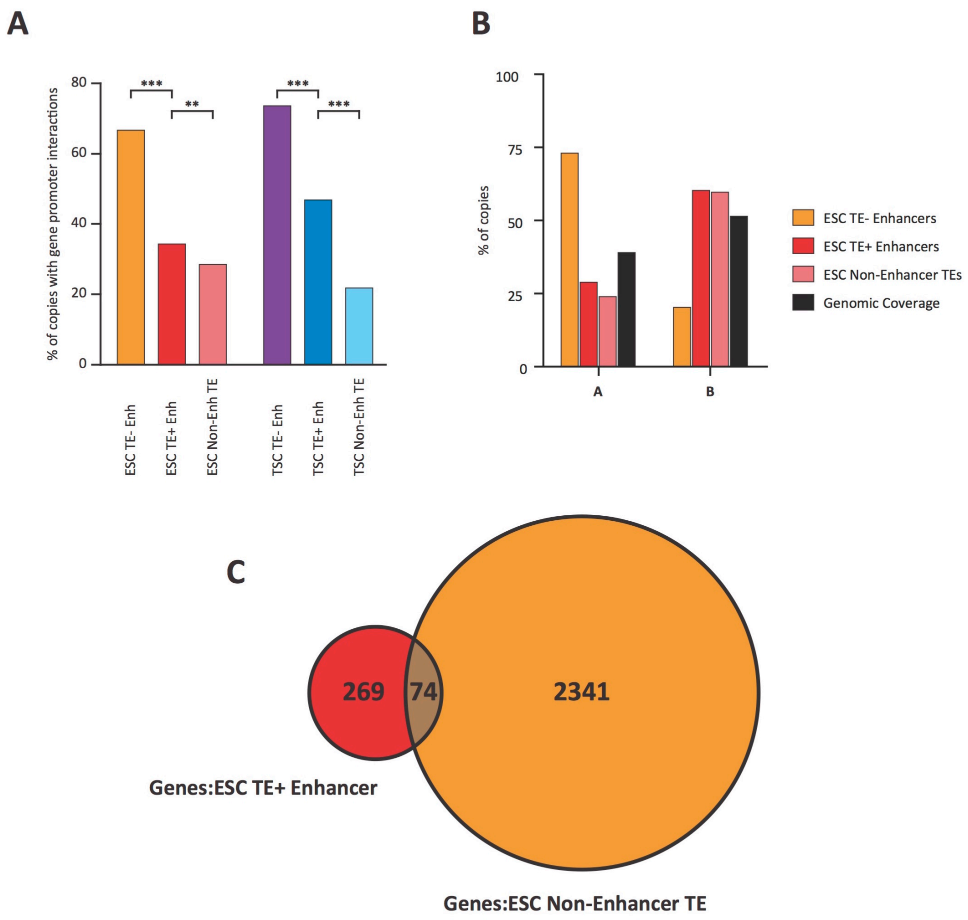


**Supplementary Figure 4 – Spatial arrangement of TEs and enhancers.** A) Proportion of elements interacting with at least one gene promoter in PCHi-C data from ESCs or TSCs (**p<0.01, ***p<0.001, proportions test with Benjamini-Hochberg correction). B) Proportion of elements within A (gene-rich, active) or B (gene-poor, inactive) spatial compartments, with the overall genomic coverage of each compartment shown for comparison. ESC compartment data are from Bonev et al. 2017. C) Venn diagram of genes contacted by TE+ or TE- enhancers in ESCs, showing only a small overlap between the two. Genes in common between the two groups were not considered for downstream analyses.

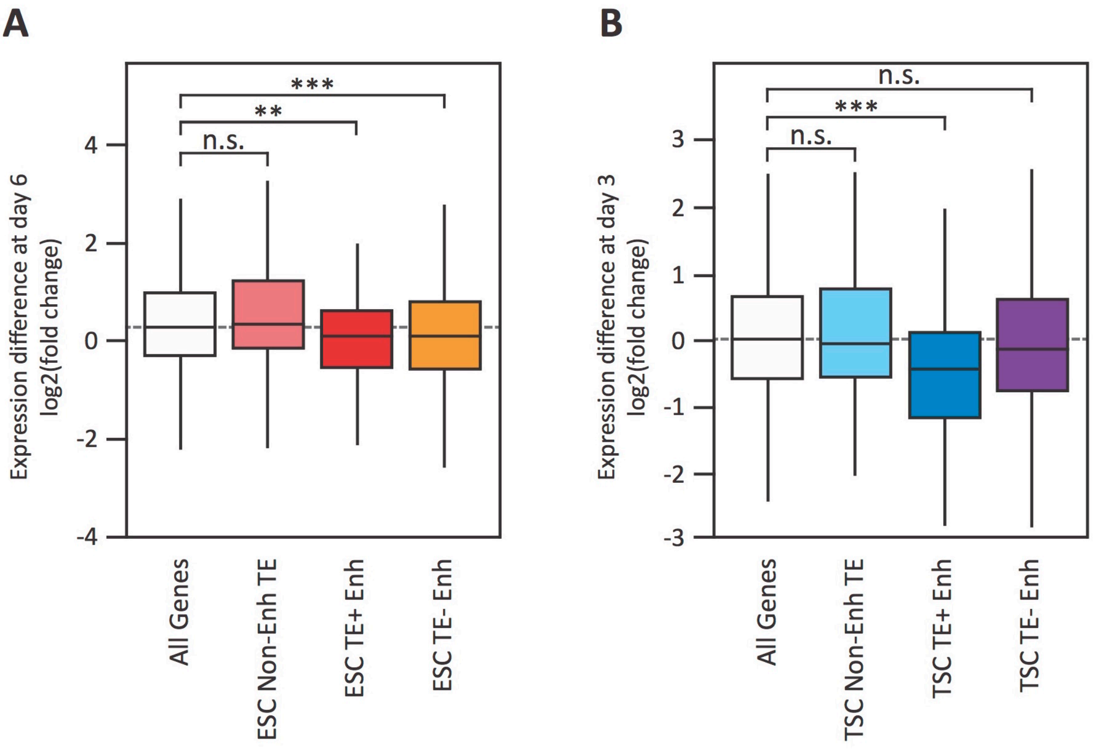


**Supplementary Figure 5 – Gene expression changes during early differentiation.** RNA-seq data from differentiating ESCs (A; Hon et al. 2014) and TSCs (B; Latos et al. 2015), showing fold change in expression of genes interacting with each of the groups indicated in Figure 3A (** $p<0.01$, *** $p<0.001$, ANOVA with Holm-Sidak's test).

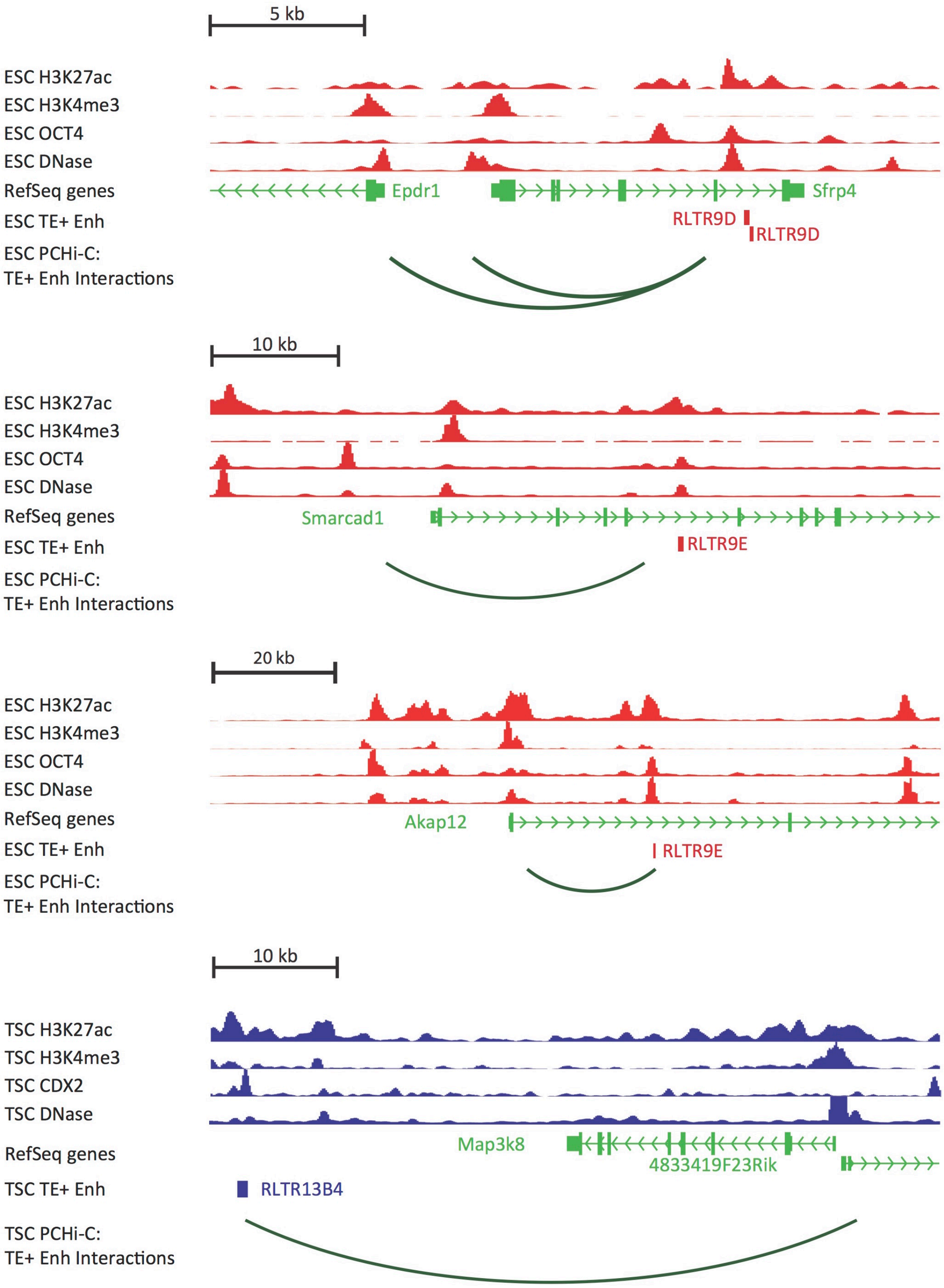


**Supplementary Figure 6 – Additional profiles of candidate TE+ enhancers.** Genome browser snapshots of TEs selected for genetic editing in ESCs or TSCs.

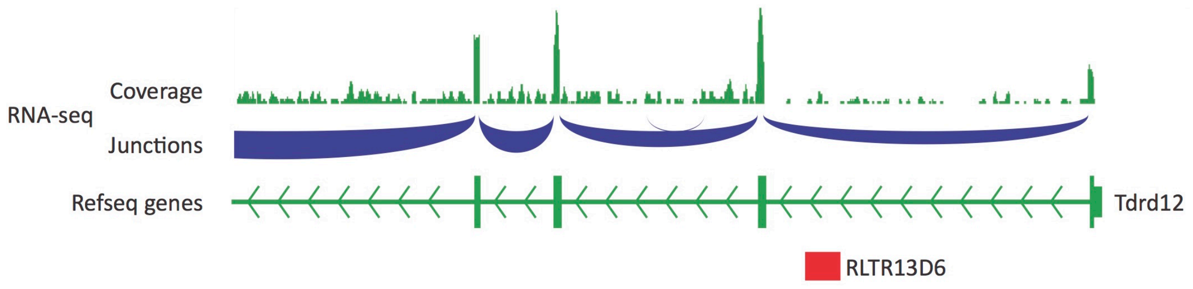


**Supplementary Figure 7 – *Tdrd12* transcriptional profile.** IGV genome browser snapshot our RNA-seq data from wildtype ESCs at the *Tdrd12* gene. Read coverage and splice junction information provide no evidence for use of the intronic RLTR13D6 element as an alternative promoter.

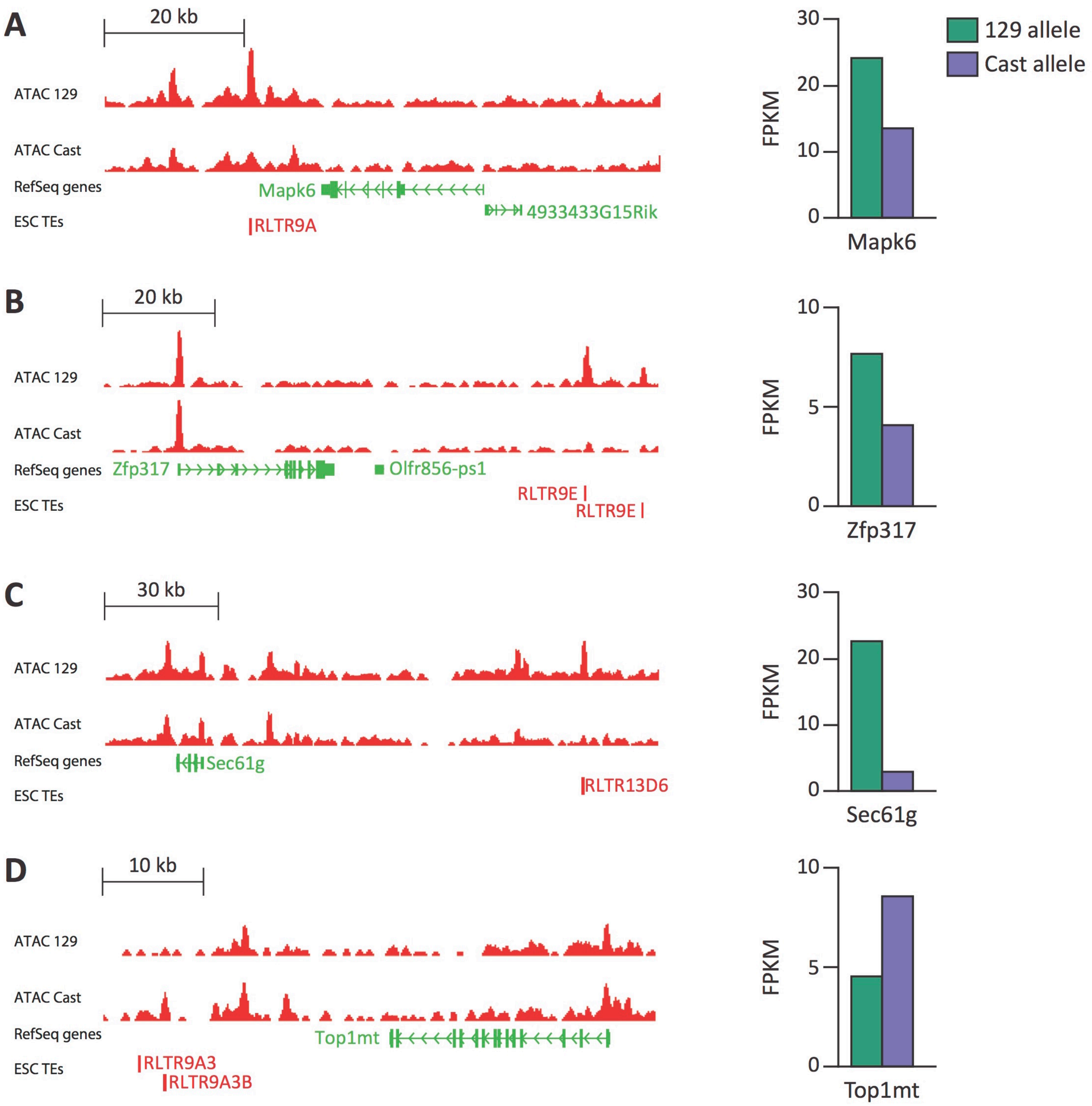


**Supplementary Figure 8 – Identification of putative TE+ enhancers based on allele-specific data.** ATAC-seq data (left) from a 129 x Cast ESC line shows examples of RLTR13D6 or RLTR9 elements with allele-specific chromatin accessibility in either the 129 (A-C) or Cast (D) allele. Allele-specific RNA-seq data from the same cell line (right) suggest a association between TE chromatin accessibility and expression of nearby genes for these examples.

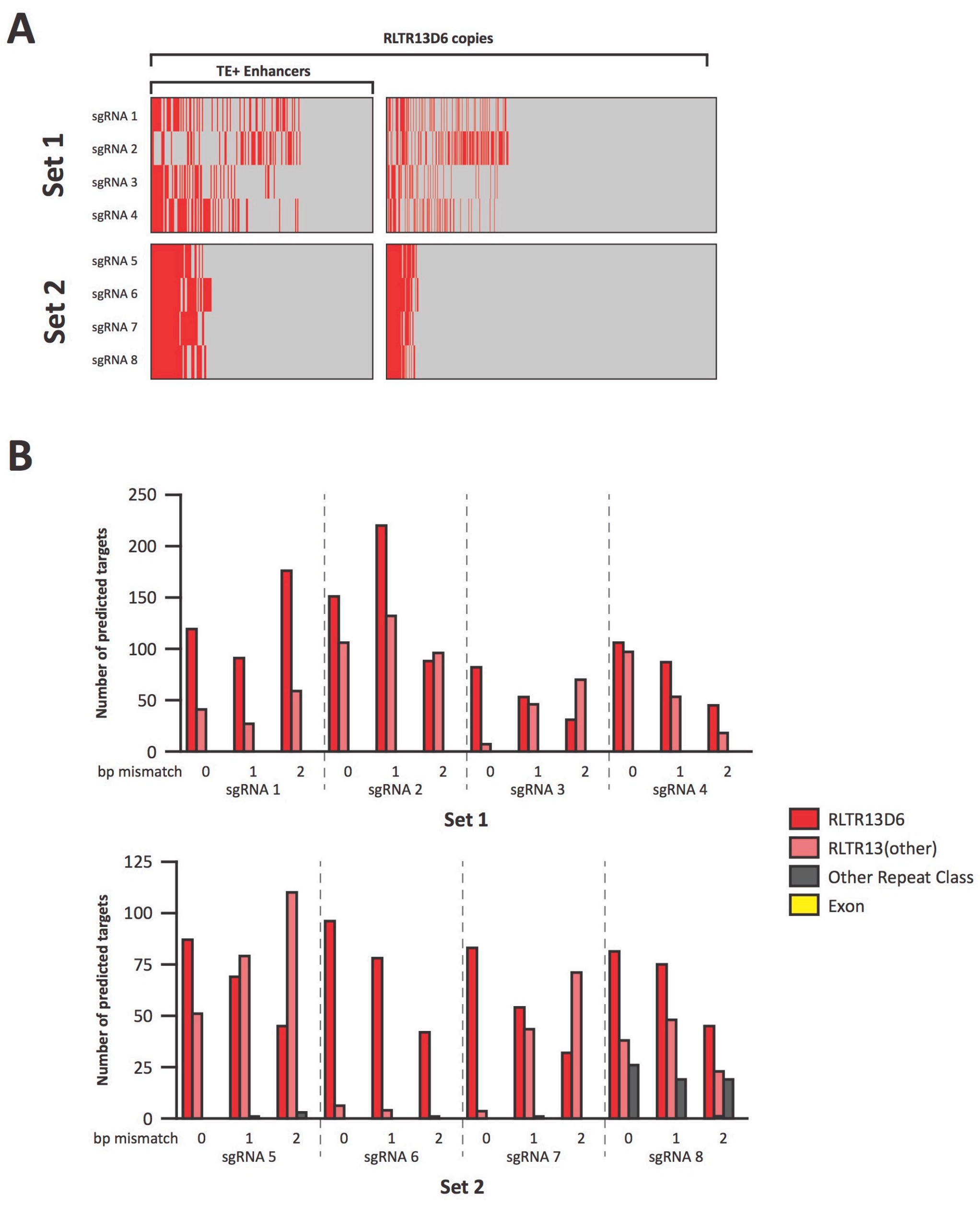


**Supplementary Figure 9 – Target predictions for sgRNAs used for CRISPRi.** A) RLTR13D6 copies predicted to be targeted (in red) by each of the sgRNAs in set 1 or set 2, with no mismatches. B) Off-target predictions for each of the sgRNAs, with either 0, 1 or 2 bp mismatch. Most off-target loci are at related TE subfamilies, and no exons are predicted to be targeted by any of the sgRNAs. Predictions are from Cas-OFFinder.

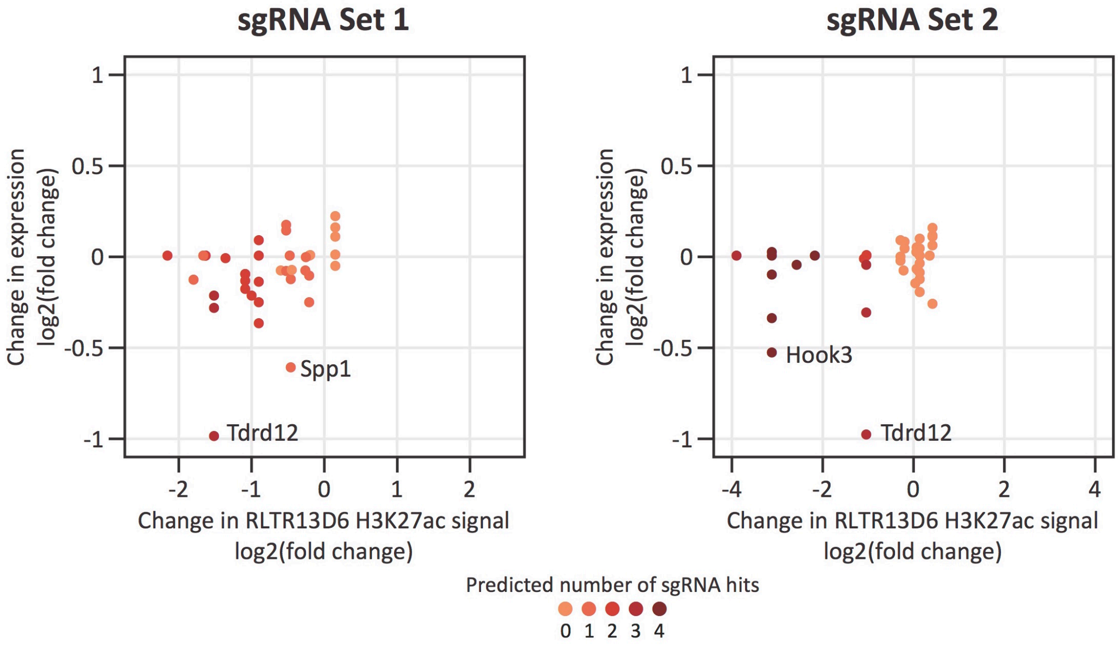


**Supplementary Figure 10 – Effects of CRISPRi on H3K27ac levels and gene expression.** Expression changes of genes interacting with H3K27ac-enriched RLTR13D6 elements are shown relative to the changes in H3K27ac levels at these TEs. *Tdrd12*, *Spp1* and *Hook3* are highlighted as the only genes with significantly differential expression upon CRISPRi – note that other genes with similar changes in H3K27ac levels (at the interacting RLTR13D6 element) remain unchanged.
